# Heterogeneous transcriptome response to DNA damage at single cell resolution

**DOI:** 10.1101/737130

**Authors:** Sung Rye Park, Sim Namkoong, Zac Zezhi Zhang, Leon Friesen, Yu-Chih Chen, Euisik Yoon, Chang H. Kim, Hojoong Kwak, Hyun Min Kang, Jun Hee Lee

## Abstract

Cancer cells often heterogeneously respond to genotoxic chemotherapy, leading to fractional killing and chemoresistance^1, 2^, which remain as the major obstacles in cancer treatment. It is widely believed that DNA damage induces a uniform response in regulating transcription and that cell fate is passively determined by a threshold mechanism evaluating the level of transcriptional responses^3^. On the contrary to this assumption, here we show that a surprisingly high level of heterogeneity exists in individual cell transcriptome responses to DNA damage, and that these transcriptome variations dictate the cell fate after DNA damage. Many DNA damage response genes, including tumor suppressor p53 targets, were exclusively expressed in only a subset of cells having specific cell fate, producing unique stress responses tailored for the fate that the cells are committed to. For instance, *CDKN1A*, the best known p53 target inhibiting cell cycle, was specifically expressed in a subset of cells undergoing cell cycle checkpoint, while other pro-apoptotic p53 targets were expressed only in cells undergoing apoptosis. A small group of cells exhibited neither checkpoint nor apoptotic responses, but produced a unique transcriptional program that conferred strong chemoresistance to the cells. The heterogeneous transcriptome response to DNA damage was also observed at the protein level in flow cytometry. Our results demonstrate that cell fate heterogeneity after DNA damage is mediated by distinct transcriptional programs generating fate-specific gene expression landscapes. This finding provides an important insight into understanding heterogeneous chemotherapy responses of cancer cells.

---

Upon genotoxic stresses, cells respond in various ways to minimize the damage consequences^4, 5^. Although individual cells respond differently to DNA damage, transcriptome level-studies of DNA damage response have been conducted only in bulk thus far^6^. Consequently, it is unknown whether the cells with different fates have continuous (Fig. 1a, upper) or distinct (Fig. 1a, lower) transcriptomic landscape. In the latter case, it could be questioned about how many distinct cell fates are possible in terms of transcriptomic phenotypes (Fig. 1b). It is also unknown whether a specific DNA damage response gene is upregulated in the whole population (Fig. 1c, upper) or in a specific subset of the population (Fig. 1c, lower), and whether different DNA damage-responsive genes are co-expressed (Fig. 1d, upper) or expressed in a mutually exclusive way in different groups of cells (Fig. 1d, lower).

**Fig. 1.**
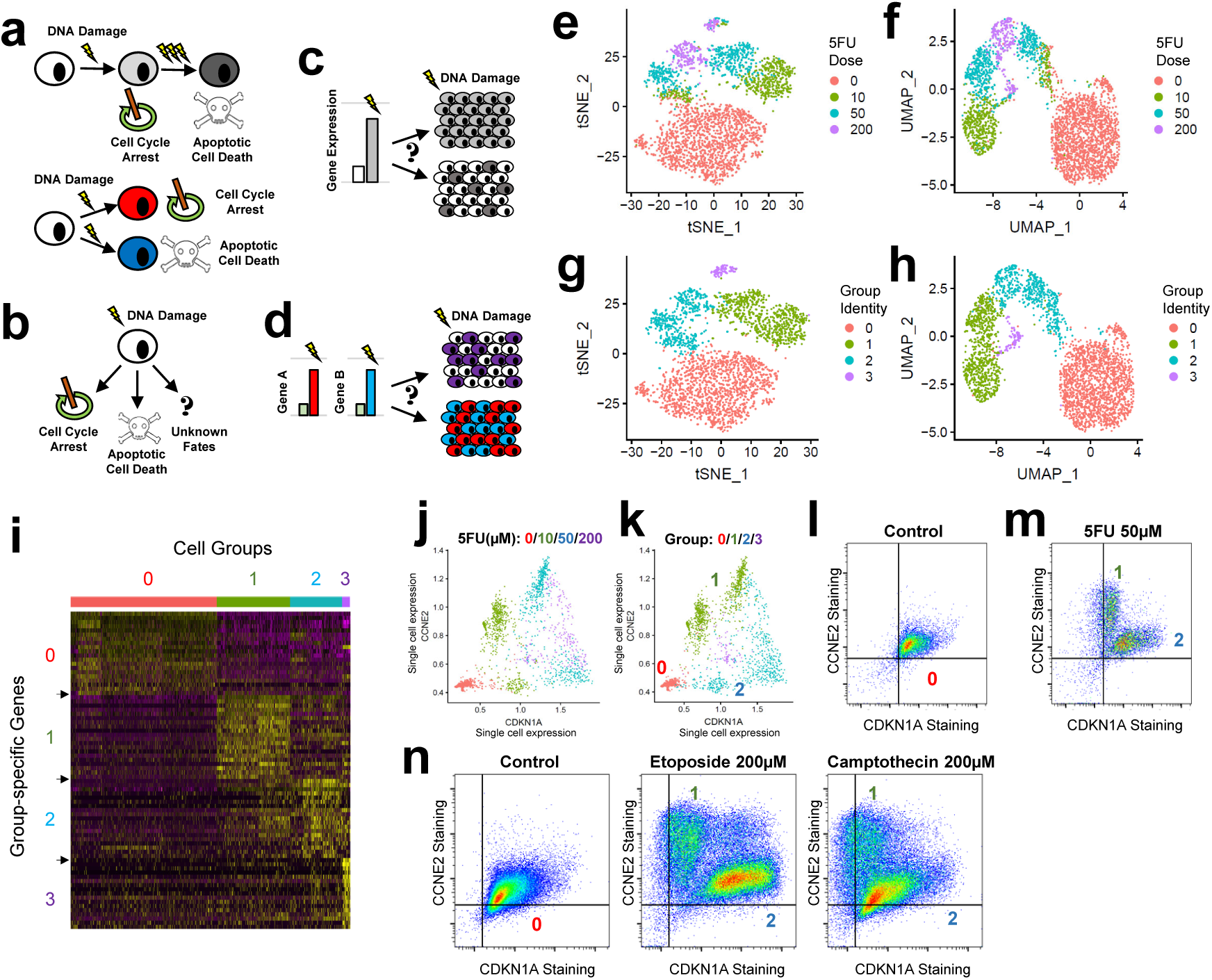
5FU treatment induces heterogeneous transcriptome response to DNA damage. **a-d**, Diagrams depicting main questions in this paper. **e-h**, t-SNE and UMAP manifolds colored with 5FU dose (**e, f**) and group identity (**g, h**) information. Group identity was determined through high-dimensional clustering of scRNA-seq data. Untreated cells fell into a single group (group 0), while 5FU-treated cells were clustered in three different groups (groups 1-3). **i**, Heat map analysis depicting expression of top 20 markers for each of the four groups identified at the top. **j, k**, Scatterplot of indicated mRNA expression in single cells, calculated from Markov affinity-based graph imputation of cells (MAGIC)^17^. Each dot represents an individual cell colored with dose of 5FU treatment (**j**) or its group identity (**k**). **l-n**, Flow cytometry analysis of indicated protein abundance in single cells. Horizontal and vertical lines indicate background fluorescence levels determined from unstained cells.

To address these questions, we performed Drop-seq^7^ and determined a total of 10,421 single cell transcriptome profiles from three different colon cancer cell lines: RKO, HCT116 and SW480. These cells were either untreated or treated with different doses of a genotoxic chemical 5-fluorouracil (5FU), in 10 independent Drop-seq experiments (Extended Data Fig. 1a, b). In principal component analysis (PCA), t-distributed stochastic neighbor embedding (t-SNE), and uniform manifold approximation and projection (UMAP)^8^, different cell lines displayed distinct transcriptomic phenotypes (Extended Data Fig. 1c). For all of these cell lines, 5FU-treated cells were clustered in locations that were distinct from untreated cells (Extended Data Fig. 1d, e), indicating that the 5FU-induced DNA damage altered single cell transcriptomes. Indeed, 5FU regulated formerly known DNA damage response genes, although the level and frequency of regulation were different across the cell lines (Extended Data Fig. 1f-h).

Using the RKO dataset, which exhibited the most robust DNA damage response from our dataset (Extended Data Fig. 1f-h), we explored the heterogeneity of single cell transcriptome profiles with high-dimensional clustering. We identified four major clusters of cells (Fig. 1e, f and Extended Data Fig. 2a), where one cluster (group 0; n = 1,597) mainly consists of untreated samples, while the other three clusters (groups 1, 2 and 3; n = 800, 571 and 85, respectively) correspond to 5FU-treated samples (Fig. 1g, h). Each of the three 5FU-treated clusters (groups 1-3) was found in all doses and batches (Fig. 1g, h and Extended Data Fig. 2b-e), indicating that these clustering results are not simply based on dose- or batch-specific effects.

The top 30 genes specifically expressed in each group were isolated through differential expression analysis (Extended Data Table 1). Each of the 5FU-treated groups has a unique subset of specific genes, which are strongly upregulated in the corresponding group but not the other groups (Fig. 1i). For instance, *CCNE2* and *CDKN1A* are among the top differentially regulated genes between the groups 1 and 2 (Extended Data Table 1). Although both genes were formerly known to be induced upon 5FU treatment in a dose-dependent manner (Fig. 1j)^9^, strong induction of *CCNE2* and *CDKN1A* was only observed in groups 1 and 2, respectively (Fig. 1k). Congruent with the scRNA-seq data, flow cytometry analyses clearly demonstrated that 5FU treatment provoked a single homogenous untreated population of CCNE2-low CDKN1A-low cells (Fig. 1l) to differentiate into two distinct subpopulations: CCNE2-high CDKN1A-low group 1-like cells, and CCNE2-low CDKN1A-high group 2-like cells (Fig. 1m). Similar pattern was also observed when the cells were treated with different doses of 5FU (Extended Data Fig. 3a) or when another group 2-specific gene MDM2 was used instead of CDKN1A (Extended Data Fig. 3b, c). Importantly, treatment with camptothecin and etoposide, two genotoxic drugs unrelated to 5FU, also induced the emergence of the CCNE2-high CDKN1A-low group 1 and the CCNE2-low CDKN1A-high group 2 (Fig. 1n). These results indicate that the heterogeneous single cell response to DNA damage is a general response to different types of genotoxic drugs.

According to the gene ontology database of biological pathways (GO-BP) analyses^10^ of the top 30 group-specific genes, group 1, the largest 5FU-treated group, expressed genes involved in the apoptotic pathway (Fig. 2a). In contrast, group 2, the second largest 5FU-treated group, expressed DNA damage-induced cell cycle checkpoint genes (Fig. 2a). Group 3, the group with fewest number of cells, was enriched with genes mediating stress responses (Fig. 2a). These analyses suggest that cells in groups 1, 2 and 3 assume different cell fate responses after 5FU treatment: apoptosis, cell cycle checkpoint and stress response, respectively (Fig. 2b).

**Fig. 2.**
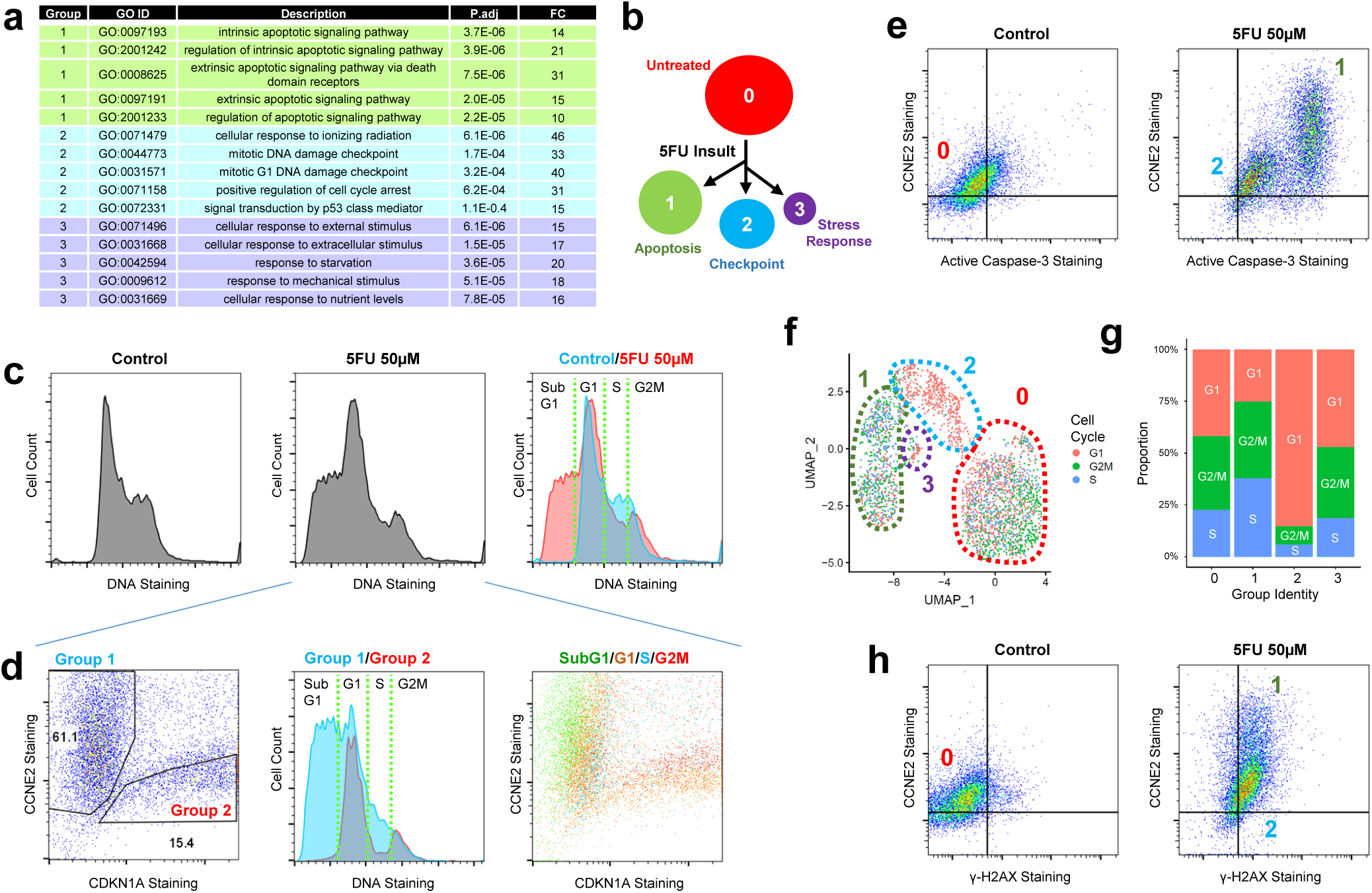
Single cell transcriptome profile dictates the fate of cells after DNA damage. **a**, Gene ontology-biological pathways (GO-BP) enrichment analysis of the top 30 markers for each of the three 5FU-treated groups. Top 5 GO-BP terms, ordered by adjusted P values (P.adj), were summarized in the table. FC, fold enrichment. **b**, Schematic model depicting 5FU-induced transcriptomic responses and their relationship to cell fate. **c**, DNA content analysis of control (left) and 5FU-treated (center) cells in flow cytometry. The two graphs were merged for a direct comparison (right). **d**, Using data from the 5FU-treated cells (center panel in **c**), cells in the groups 1 and 2 were separately analyzed for their DNA content. Gating scheme was shown in the scatterplot of CCNE2 and CDKN1A (left). DNA content of cells were analyzed in each gate representing the group 1 (blue, center) and 2 (red, center). Stages of cell cycle were estimated through the DNA content analysis (center), and used to color single cells in the scatterplot of CCNE2 and CDKN1A (right). **e, h**, Flow cytometry analysis of indicated protein expression. Horizontal and vertical lines indicate background fluorescence level determined from unstained cells. **f**, The stage of cell cycle was estimated through analyzing expression of S and G2/M phase-specific markers Approximate boundaries for each group are marked by a dotted line. **g**, Proportion of cells in each group classified into different cell cycle stages.

To further substantiate these observations, we performed the DNA content analysis through flow cytometry. As predicted, 5FU treatment induced strong accumulation of a sub-G1 population, which is suggestive of cell death (Fig. 2c). The 5FU-treated cell population was partitioned into the CCNE2-high CDKN1A-low group 1 and the CCNE2-low CDKN1A-high group 2, based on their flow cytometry profiles (Fig. 2d, left). These two groups of cells exhibit different forward versus side scatter analyses values, indicating that their cell size and internal complexity are different from each other (Extended Data Fig. 3d). Roughly half of the group 1 cells appear as sub-G1 (Fig. 2d, center), indicating that this group contain many dying cells. Analyses with active caspase-3 further confirmed that all cells in the CCNE2-high group 1 indeed have activated apoptotic caspase cascade (Fig. 2e). In contrast, group 2 did not have sub-G1 cells (Fig. 2d, center) and expressed relatively low levels of active caspase 3 (Fig. 2e), indicating that they are protected from apoptosis. Group 2 cells instead exhibited strong G1 arrest in both flow cytometry (Fig. 2d, center) and scRNA-seq (Fig. 2f, g) analyses. These results demonstrate that the group 1 cells were indeed undergoing apoptosis while the group 2 cells experience cell cycle arrest, as suggested by the gene ontology study (Fig. 2b). Although groups 1 and 2 exhibited different cell fate phenotypes, they experienced similar levels of DNA damage, as assessed by the γ-H2AX expression (Fig. 2h).

As 5FU dose becomes higher, the number of group 1 cells from the scRNA-seq dataset was strongly decreased (Extended Data Fig. 2c-e), although this pattern was not observed in flow cytometry (Extended Data Fig. 3a). It is possible that high levels of DNA damage accelerated apoptotic progression of the group 1 cells, producing apoptosis-associated RNA decay^11^. Therefore, even though the number of cells in the group 1 were high in flow cytometry (Fig. 3a), they would not be represented in the scRNA-seq data (Extended Data Fig. 2c-e).

**Fig. 3.**
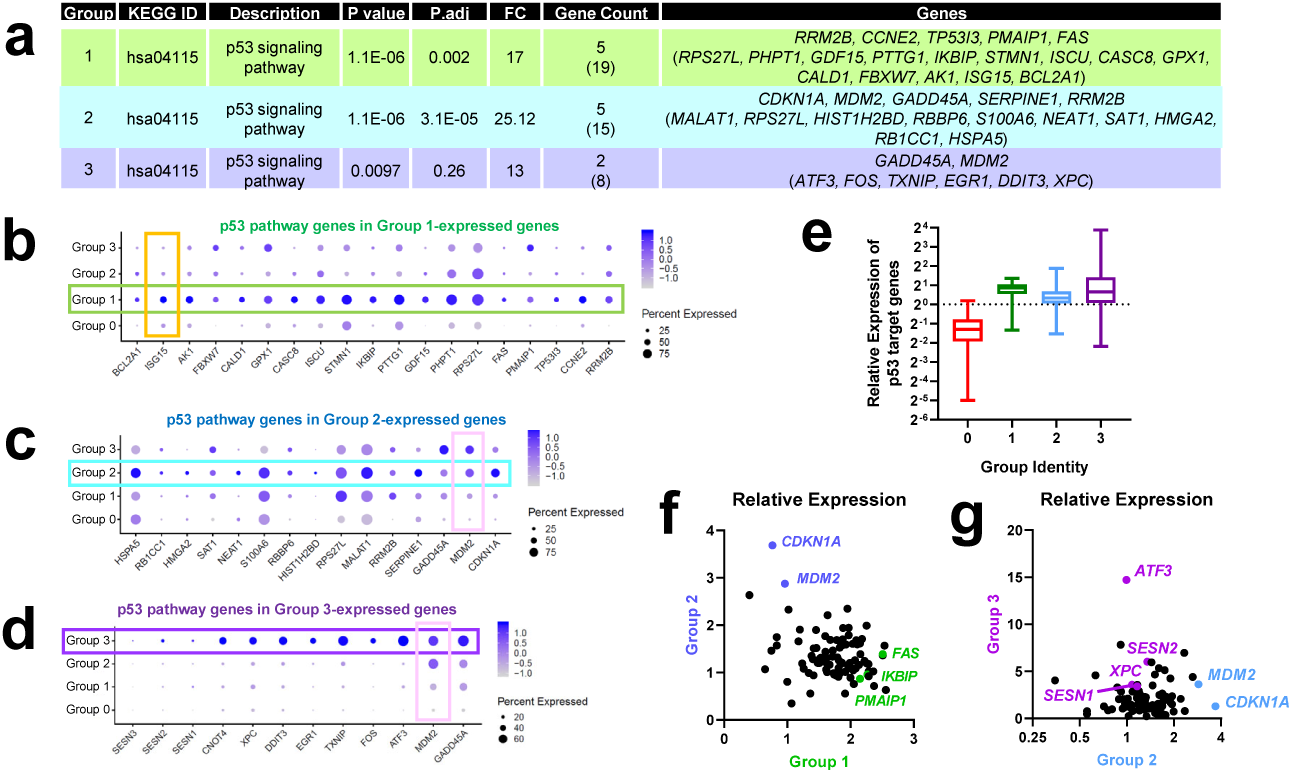
Unique expression patterns of p53 pathway genes according to the group identities of single cells. **a**, Pathway enrichment analysis using the Kyoto Encyclopedia of Genes and Genomes (KEGG) database identified the p53 pathway genes as the most important feature distinguishing the three 5FU-induced groups. Counts in parentheses are a subset of the top 30 genes whose involvement in p53 pathway was documented in the literature but was not included in the KEGG database. P.adj, adjusted P value; FC, fold enrichment. **b-d**, Dot plot of the p53 pathway genes that were in the top 30 markers for groups 1 (**b**), 2 (**c**) and 3 (**d**). The size of the dot reflects the percentage of cells expressing the markers, while the color encodes average expression levels across all cells within the group (blue is high). *CNOT4* and *SESNs* are not in the top 30 markers but were significantly upregulated in the group 3. **e-g**, Average expression level of individual p53 target genes in each group, normalized by their averaged expression in all cells, are presented in box plot (**e**) and correlation scatterplots (**f, g**). Total 89 p53 target genes were analyzed. Each dot in the scatterplots represents an individual p53 target gene.

The group 3 is a unique group of cells that is small but consistently detected in all experiments regardless of 5FU doses. Unlike group 1 cells, the abundance of group 3 cells did not decrease after high-dose 5FU treatments (Extended Data Fig. 2c-e). Also unlike group 2 cells, the group 3 cells did not undergo cell cycle arrest (Fig. 2f, g); therefore, group 3 cells seem to have evaded both apoptosis and cell cycle checkpoint responses. As the group 3 cells express high levels of genes mediating stress response, such as *ATF3, FOS* and *DDIT3* (Extended Data Fig. 4a, b), it is likely that they represent a novel fate of chemoresistance. Group 3-like cells were also identified from flow cytometry as CCNE2-low and ATF3/FOS-high cells (Extended Data Fig. 4c-f).

Consistent with the notion that p53 pathway is central to cellular DNA damage response, all 5FU-treated groups identified the p53 signaling pathway as the top Kyoto Encyclopedia of Genes and Genomes (KEGG)^12^ pathway enriched in each group (Fig. 3a). However, intriguingly, a vast majority (93%) of these marker genes were exclusively found in a single group, and even the remaining three genes (*MDM2, GADD45A* and *RRM2B*; 7%) were found only in two groups but not in the third (Fig. 3b-d and Extended Data Fig. 5). For instance, *MDM2*, a well-characterized negative feedback regulator of p53, was very highly expressed in the groups 2 and 3, but not in the group 1 (Fig. 3c, d; pink boxes). In contrast, *ISG15*, a recently identified positive feedback regulator of p53^13^, was highly expressed only in the group 1, but not in the groups 2 and 3 (Fig. 3b; yellow box). It is possible that ISG15-mediated positive feedback in group 1 cells allowed for sustained p53 activation^13^, leading to higher p53 activity and apoptotic cell death^14^. MDM2-mediated negative feedback in group 2 and 3 cells may have produced pulse responses in p53 activities^15^, inducing non-apoptotic consequences of limited p53 activation^14^.

Using a recently assembled list of p53 target genes^16^, we systematically investigated the 5FU-dependent p53 target activation in different groups of cells. For most of the p53 targets, 5FU-induced expression was higher in the groups 1 and 3 than in the group 2 (Fig. 3e). For instance, many pro-apoptotic p53 target genes, such as *PMAIP1, FAS* and *IKBIP*, were highly expressed in the group 1 (Fig. 3f and Extended Data 6), and p53 target genes important for stress resistance, such as *ATF3, XPC* and *SESN1-3*, were most distinctly expressed in the group 3 (Fig. 3g and Extended Data 5c, f). Only a small number of genes, such as *CDKN1A* and *MDM2*, were expressed highly in group 2, compared to the other groups (Fig. 3f, g and Extended Data 6). Therefore, even though all 5FU-treated groups generally up-regulated p53 pathway genes, they induced a different subset of genes.

These results provide clear and convincing answers to all the questions we initially raised (Fig. 1a-d). DNA damage induces distinct transcriptional programs leading to diversified cell fates (Fig. 1a, lower path), which includes, in addition to classical apoptosis and cell cycle checkpoints fates, a novel chemoresistant fate (Fig. 1b). Many DNA damage-induced genes, including tumor suppressor p53 targets, are differentially expressed among the different groups of cells (Fig. 1c, lower path), which is often mutually exclusive to each other (Fig. 1d, lower path). Interestingly, the DNA damage-induced transcriptome differentiation phenotype, strongly manifested in the RKO cell line, was less pronouncedly observed in other cell lines (Fig. 4). Although RKO cells display clear distinction between groups 1, 2 and 3 (Fig. 4a-f), HCT116 cells showed separation of only groups 1 and 2 (Fig. 4g-l). SW480 cells, whose DNA damage response is compromised by p53 mutation, did not exhibit significant transcriptome differentiation, and they only exhibited dose- and batch-specific transcriptome phenotypes (Fig. 4m-r). These results suggest that an intact DNA damage response is important for producing heterogeneity in transcriptome responses.

**Fig. 4.**
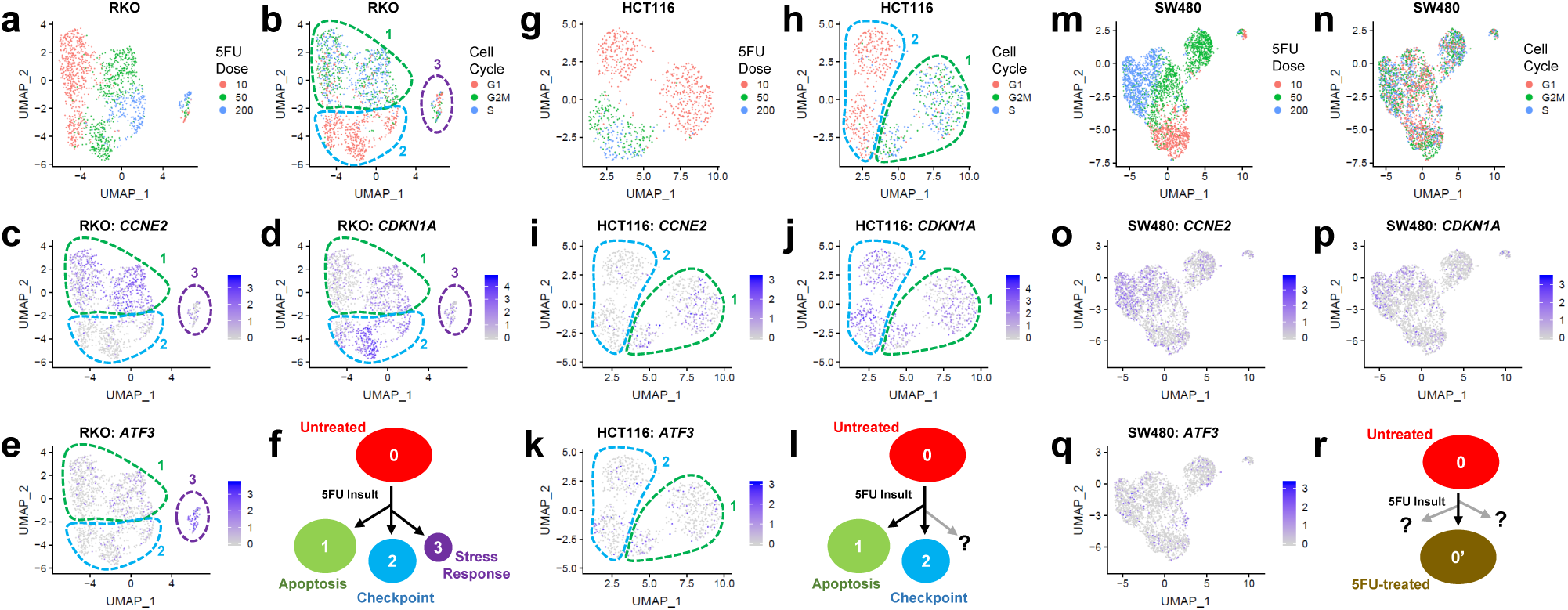
Patterns of single cell DNA damage response across different colon cancer cell lines. UMAP manifolds and feature plots of 5FU-treated RKO (**a-e**), HCT116 (**g-k**) and SW480 (**m-q**) cells. UMAP manifolds were colored by 5FU treatment dose (**a, g, m**) or estimated cell cycle phase (**b, h, n**). Feature plots of group-specific genes were generated for *CCNE2* (group 1; **c, i, o**), *CDKN1A* (group 2; **d, j, p**) and *ATF3* (group 3; **e, k, q**). Models of 5FU-induced transcriptome differentiation in RKO (**f**), HCT116 (**l**) and SW480 (**r**) cells are illustrated.

In summary, we provide the first snapshot of how individual cells shape their transcriptome in response to DNA damage. This work unveils new information about how different genes are expressed in different subgroups of cells sharing a specialized cell fate. In addition, by revealing a cell fate-specific transcriptome patterns, we open a new avenue for future studies to understand heterogeneous cancer cell responses to genotoxic chemotherapy, such as fractional killing and chemoresistant tumor recurrence.

## Acknowledgements

We thank Dr. Y. Shah for colon cancer cell culture, Drs. M. Kim and C. S. Cho for advices, B. Gu, I. A. Semple and B. Kim for technical assistance, A. H. Kowalsky for comments, and Santa Cruz Biotechnology for antibodies. We also appreciate Drs. K. R. Spindler (Univ. of Michigan) and Y. Xi (Univ. of Texas at Austin) for their intellectual inputs. This work was supported by the NIH grants (R01DK102850 and R01DK114131 to J.H.L., U01HL137182 to H.M.K., and P30AG024824, P30DK034933, P30DK089503, and P30CA046592), the Chan Zuckerberg Initiative (to H.M.K.), the MCubed initiative (to J.H.L., H.M.K. and C.H.K.), Dean’s Organogenesis Fellowship (to S.N.), and the AASLD Pilot Research Award (to J.H.L. and H.M.K.).

## Author contributions

S.R.P. performed all experiments. S.R.P. and H.M.K. conducted computational analysis of Drop-seq data. S.N., Y.C.C., E.Y. and H.K. contributed to establishing Drop-seq experiments. Z.Z.Z. contributed to statistical analysis. L.F. and C.H.K. contributed to the analysis of flow cytometry data. H.M.K. and J.H.L. conceived and directed the project. S.R.P., H.M.K. and J.H.L. designed experiments, analyzed data and wrote the manuscript. All authors revised or commented on the manuscript and approved the final version.

## Competing interests

The authors declare no competing interests.

## Methods

### Cell culture

RKO (CRL-1577), HCT116 (CCL-247) and SW480 (CCL-228) cells were obtained from the American Type Culture Collection (ATCC) and maintained in Dulbecco’s Modified Eagle Medium (DMEM; Fisher Scientific, 11965-092, Gibco) with 10% fetal bovine serum (FBS, Sigma, F4135) and 1% penicillin-streptomycin (Fisher Scientific, 15140122) at 37°C and 5% CO_2_. Upon receipt, each cell line was subcultured for fewer than 6 months prior to initiation of the described experiments. All cell lines were negative for Mycoplasma contamination in PCR-based analysis using the following primers: F: 5’- GTGGGGAGCAAA(C/T)AGGATTAGA-3’, R: 5’-GGCATGATGATTTGACGTC(A/G)T-3’. In addition, cell lines were again authenticated through sequence comparison with COSMIC database (RKO and HCT116)^18^ and through profiling 15 autosomal short tandem repeat (STR) loci and the gender identity locus (SW480, Genetica Cell Line Testing).

### DNA damage treatment

Cells were seeded at 100 cells/mL in DMEM with FBS. When the cells reached 70% confluence, DNA damage was induced by treatments with 5-fluorouracil (10µM, 50µM and 200µM, as indicated) for Drop-seq and flow cytometry experiments. Etoposide (200µM) or camptothecin (200µM) were also treated for 24 hours for flow cytometry experiments. Control cells were harvested before the DNA damage treatment for all experiments.

For Drop-seq, individual cell lines with indicated treatment were harvested after TrypLE Express digestion (ThermoFisher, 12605010, Gibco). After centrifugation at 3500 rpm for 5 min, cells were washed in Dulbecco’s phosphate-buffered saline (DPBS; Fisher Scientific, 14190250) and centrifuged again. The cells were then resuspended in DPBS containing 0.1% bovine serum albumins (Sigma, A8806). Then the cells were filtered through a 40-micron cell strainer (VWR 21008949) and diluted to the concentration of 100,000 cell/ml. Cells from three different cell lines were then mixed with each other right before being subjected to the Drop-seq experiment. The cells for flow cytometry were harvested using the same protocol and diluted into DPBS before the fixation.

### Drop-seq and library preparation

We performed Drop-seq through the described protocol (Macosko et al., 2015). We mixed equal numbers of cells from three different cell lines (approximately 40,000 cells for each cell line) for one loading of 120,000 cells for Drop-seq. Briefly, three pump-controlled syringes with cell suspension (100,000 cells/ml, total 1.2 ml per run), barcoded beads (Chemgenes, MACOSKO-2011-10) in lysis buffer (400mM Tris pH 7.5, 40mM EDTA, 12% Ficoll PM-400, 0.4% Sarkosyl and 100mM DTT; 100,000 beads/ml, total 1.2 ml per run), and droplet generation oil (Bio-rad, 1864006; 7 ml per run), were connected to a microfluidics device (FlowJEM) under microscope supervision. During droplet generation, we set the cell and bead flow speed at 2,000 µl/hr, and the oil speed at 7,500 µl/hr. The droplets were collected into 50 ml falcon tubes during average time of 25 to 30 mins. Following droplet breakage, the total beads were collected and washed in 6X SSC (diluted from 20X SSC, Invitrogen, #15557044). Excess bead primers without RNA molecules were removed by the treatment of Exonuclease I (NEB, NEBM0293S). Then, we performed first strand cDNA synthesis using Template Switch Oligo (TSO) on beads. PCR cycling followed the original Drop-seq protocol (Macosko et al., 2015). The resultant PCR products were extracted using AMPure XP beads (Beckman Coulter, A63881). After this step, the number of Single-cell Transcriptomes Attached to MicroParticles (STAMPs) per run was estimated by the PCR product concentration. We used the samples that contained more than 400 STAMPs for the secondary PCR using the TSO PCR primer in order to amplify the cDNA. The resulting full-length enriched cDNA library was processed into the sequencing library using the Nextera XT DNA Library Preparation Kit (Illumina, FC-131-1096). The quality of the libraries was inspected by size and concentration by agarose gel electrophoresis before pooling the libraries. They were evaluated again with the BioAnalyzer in the UM Sequencing Core. A total of 10 sets of cDNA libraries from Drop-seq runs were analyzed. The library pools with the average size of 450 bp were sequenced using Illumina HiSeq-4000 High-Output, with asymmetrical reads of 26 bp × 110 bp.

### Flow cytometry

Cells were prepared as described above in DNA damage treatment section. Cells were washed with DPBS twice and resuspended in 500 µl DPBS. Then, the cells were processed according to the Intracellular Flow Cytometry Staining Protocol provided by Biolegend^19^ with minor modifications. In brief, cells were fixed in Fixation Buffer (BioLegend, 420801) for 20 minutes at room temperature in the dark and washed in Cell Staining Buffer (BioLegend Cat. No.420201) twice. Cells were then washed twice with 1X Intracellular Staining Perm Wash Buffer (BioLegend Cat. No.421002) for permeabilization and resuspended for antibody incubation. The primary antibodies (1 µl/sample) were applied to the Perm/Wash buffer and incubated with cells for 1 hour at room temperature in dark. p21 (#2947), MDM2 (#86934), Cleaved caspase 3 (#9661) and p-H2AX (#2577) are from Cell Signaling, and cyclin E2 (sc- 28351), ATF-3 (sc-518032) and c-fos (sc-166940) are from Santa Cruz Biotechnology. The primary antibodies were then washed out with the Perm/Wash buffer. The fluorescence-conjugated secondary antibodies (Alexa Fluor 488: A-21206, Invitrogen; Alexa Fluor 594: A-11032, Invitrogen) were added to the Perm/Wash buffer (1 µl/sample) and incubated 1 hour at room temperature in the dark. DAPI (D21490, Invitrogen) was added to the secondary antibody solution when necessary. After final wash with Perm/Wash buffer, cells were resuspended in 500 ul of Cell Staining Buffer and examined in Bio-Rad ZE5 or Fortessa Cell Analyzers. Data analysis was done using FlowJo software.

### Drop-seq data processing and cell line de-multiplexing

We processed raw reads following the instructions described in the Drop-seq Laboratory Protocol v3 using DropSeqTools (v1.13)^7^. Reads were aligned to hg19 genome using STAR (v.2.6.0a)^20^ following the default DropSeqTools pipeline. The aligned reads were further processed using *popscle dsc-pileup* to produce the profiles of Unique Molecular Identifier (UMI) profiles and pileups of reads overlapping with polymorphic variants with minor allele frequency (MAF) > 1% from 1000 Genomes Phase 3 panel^21^. The UMI profiles were examined to produce the knee plot, which displays the empirical cumulative distribution of UMIs across all barcoded droplets (Figure S1A, center). We used a UMI count 800 as a cutoff to determine the droplets to consider for the downstream analysis. As a result, 13,801 barcoded droplets passed the UMI cutoff.

The deconvolution of cell types was performed using *popscle freemuxlet*, by clustering them into 3 multiplexed samples (with --nsample 3) using variant-overlapping reads while detecting doublets with default parameters for each batch. The genetic identities of each cluster were determined by calculating the likelihood of sequence reads given genotypes from the COSMIC database^18^ for RKO and HCT116. Because SW480 genotypes were unavailable in COSMIC, the cluster with no matching samples was assumed to be SW480. The inferred genotype likelihoods for each sample were merged across 10 batches, and the merged genotype likelihoods were used to confirm the identity of droplets using *popscle demuxlet*. 11,259 droplets were confidently inferred as singlets in both *freemuxlet* and *demuxlet* and used for the subsequent analysis. In the downstream analysis (see below), droplets containing aberrantly high content of mitochondrial mRNAs were further eliminated (Figure S1A, right); therefore, the number of droplets actually used for the analysis was reduced to 10,421 (Figure S1A, left). The digital expression matrix of these droplets was produced using *popscle plp-make-dge-matrix*, using the same gene annotation database (in GTF format) used for *DropSeqTools*.

### Cell clustering and data visualization

The digital expression matrix was processed to Seurat v3^22^ following the “standard processing workflow” in the tutorial, except for a few changes. First, we used 11% (RKO and SW480) and 21% (HCT116) as the threshold to filter out droplets with aberrantly high mitochondrial reads based on the cell line-specific inspection of QC metrics (Figure S1A, right). Second, we used 10 principal components for all manifold learning and clustering analyses to maintain consistency. Third, when finding variable genes, we chose the top genes based on average expression and dispersion (selection.method = “mean.var.plot”) to maintain consistency with analysis from Seurat v2^23^. Using the revised standard workflow, we computed principal components and 2-dimensional t-SNE^24^ and UMAP^8^ manifolds in the merged dataset, as well as dataset stratified into RKO, HCT116, and SW480 as identified by *freemuxlet* and *demuxlet*.

Clustering was performed using the shared nearest neighbor modularity optimization implemented in Seurat’s *FindClusters* function using resolution parameter as 0.2. We observed that batch effects do exist, even though they were smaller than the biological effect of separating the four primary clusters. For example, when using resolution higher than 0.2, we observed additional clusters almost exclusive for specific batches. To correct for these batch effects, we applied *CCA*^25^, *MNN*^26^, and *liger*^27^. However, in our experimental settings where technical batches and biological effects (5FU dose) can confound each other, all these methods failed to correct for (or over-corrected) batch effects to examine the biological differences more clearly than uncorrected data. Therefore, we used uncorrected data in the subsequent analyses.

Heatmaps were produced using *DoHeatmap* function in Seurat using the top 20 genes with highest log-fold changes identified from differentially expressed genes identified by *FindAllMarkers* function with default parameters. Violin plots, dot plots, and feature plots (gene expression per cell in the manifold space) were generated using *VlnPlot, DotPlot, FeaturePlot* functions, respectively. Cell cycles were inferred using *CellCycleScoring* function in Seurat using the recommended set of cell-cycle specific genes^28^.

### Pathway enrichment analysis

Differentially expressed genes by clusters were applied to clusterProfiler for pathway enrichment analysis^29^. The top 30 differentially expressed genes (based on fold-enrichment) were identified for each cluster using *FindAllMarkers* function in Seurat, and the pathway enrichment analysis were performed using *enrichGO* and *enrichKEGG* functions, respectively, with significance p-value cutoff 0.05. When the exact same list of genes was identified in multiple GO terms, only the term with the lowest p-value was presented in the table.

### Imputation of single cell expression

We performed imputation of the data using *magic* package with the default parameter^17^ to detect the relationship of genes of interest. The scatterplots and feature plots of imputed data were visualized using customized R scripts with *ggplot2*.

## Extended Data Figure Legends

**Extended Data Fig. 1.**
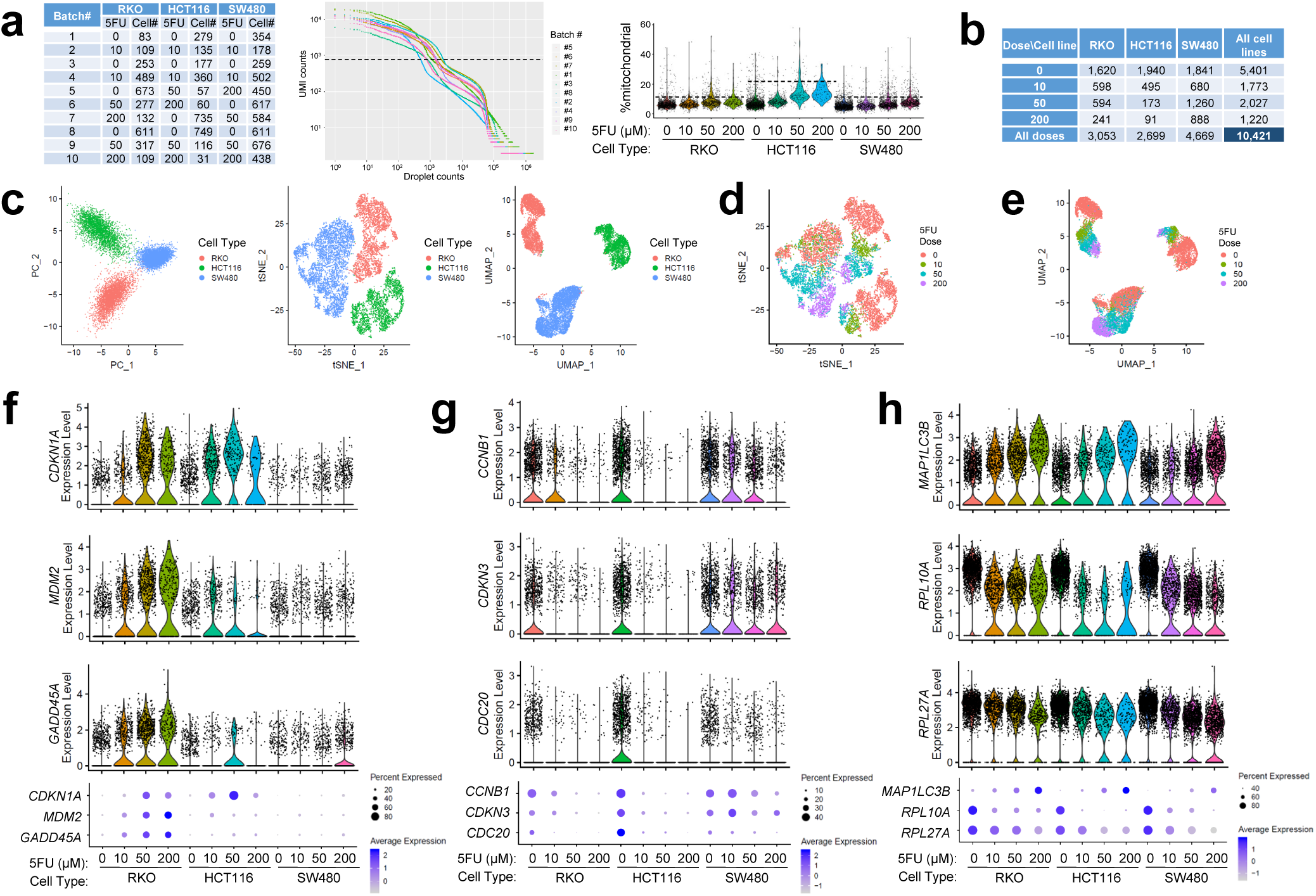
scRNA-seq captured cell line identity and DNA damage response. **a, b**, The numbers of cells captured through Drop-seq, stratified by batches (**a**, left) and 5FU treatment doses (**b**). RKO, HCT116 and SW480 cell line identities were determined through *demuxlet* and *freemuxlet* software. Knee plot (**a**, center) displays the empirical cumulative distribution of UMIs across barcoded droplets in all batches. Violin plot (**a**, right) shows the distribution of mitochondrial RNA ratio (%) across barcoded droplets in all cell lines. Cells with >800 UMIs (dotted line in the center panel of A), <11% (RKO and SW480) or <21% (HCT116) mitochondrial mRNA reads (dotted lines in the right panel of **a**), and confident inference of cell line identity were included in the analysis. **c**, PCA plot (left) and t-SNE (center) and UMAP (right) manifolds of all three cell lines colored with cell line identity. **d, e**, t-SNE (**d**) and UMAP (**e**) manifolds of all cell lines colored with 5FU dose. **f-h**, Violin plots (upper) and dot plots (lower) of the indicated DNA damage responsive genes after 5FU treatment. In the dot plots, the size of the dot reflects the percentage of cells expressing the markers, while the color encodes the average expression levels across all cells within the group (blue is high). Each cell line was analyzed separately.

**Extended Data Fig. 2.**
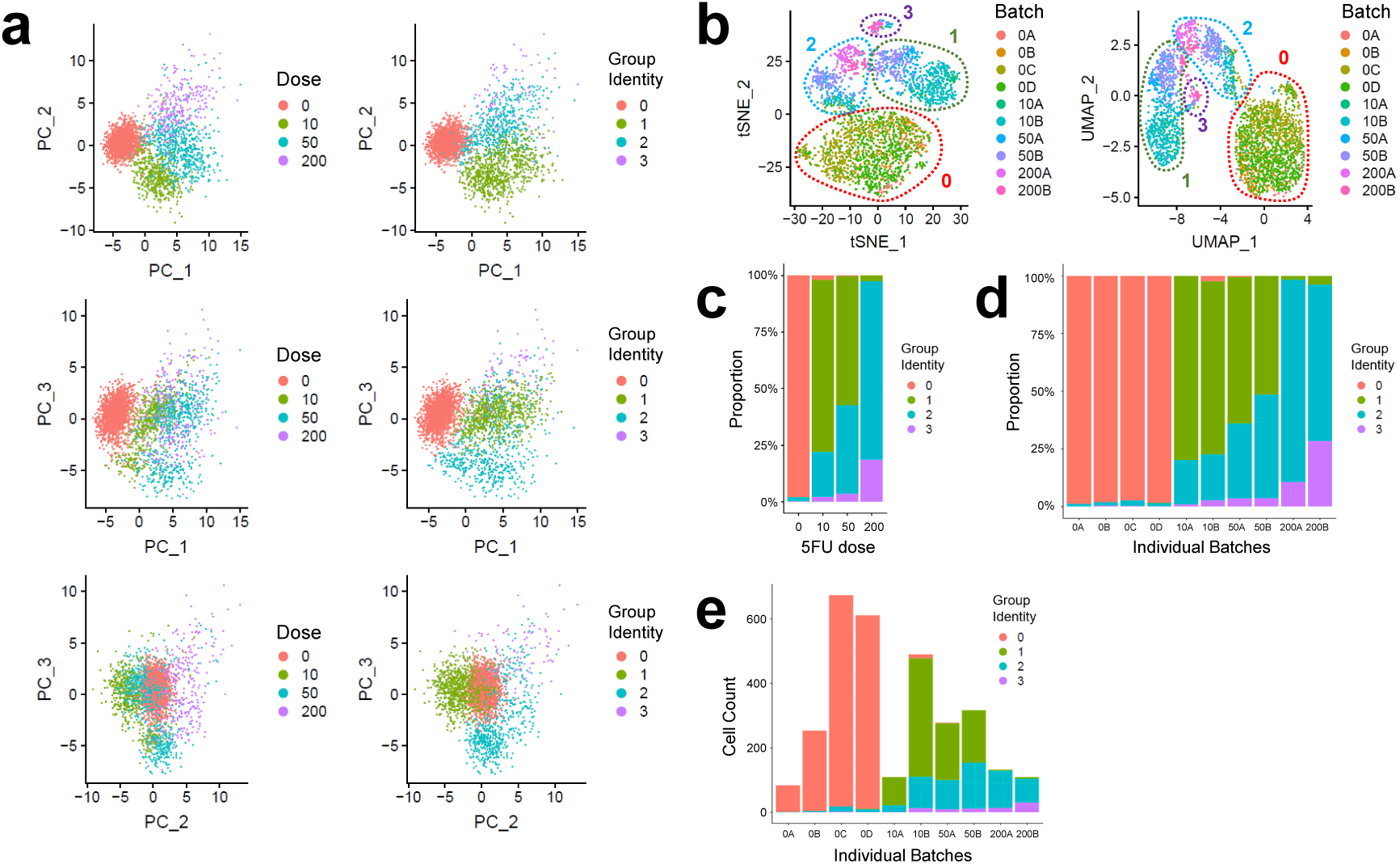
5FU-induced differentiation of single cell transcriptome in RKO cells. **a**, PCA plot of RKO cells colored with 5FU dose (left) and group identity assigned by high-dimensional clustering of scRNA-seq data (right). **b**, t-SNE and UMAP manifolds colored with batch information. Control experiments were performed in quadruplicate (0A-0D), while experiments using each 5FU dose was performed in duplicate (10A, 10B, 50A, 50B, 200A and 200B). Approximate boundaries for the groups 0, 1, 2 and 3 are marked by a dotted line. **c-e**, Distribution of cells in each dose (**c**) or batch (**d, e**) into the group identity assigned by clustering analysis.

**Extended Data Fig. 3.**
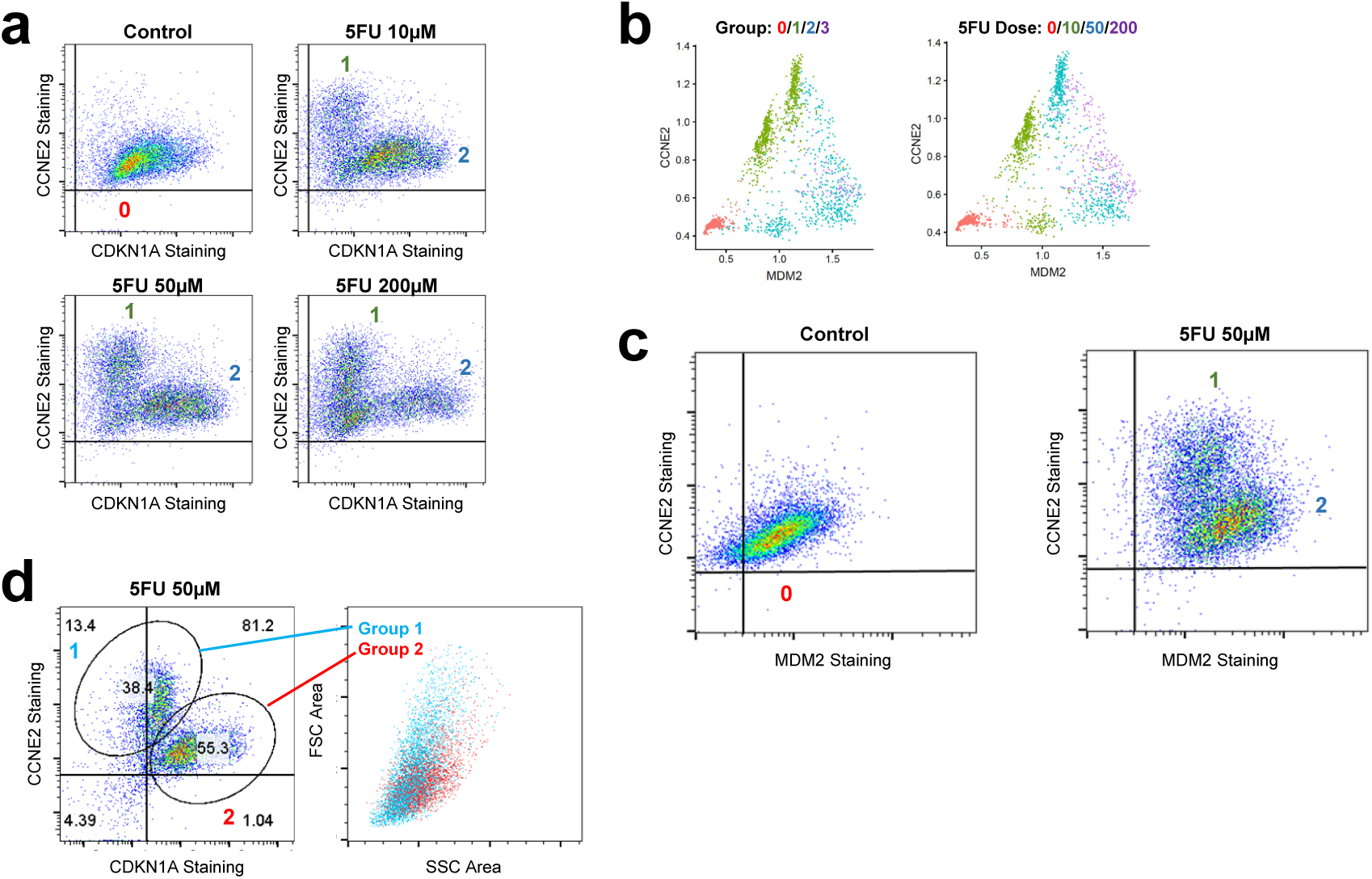
Flow cytometry confirms 5FU-induced differentiation of group 1 and group 2. **a-c**, Flow cytometry analysis of indicated protein abundance in single cells. Horizontal and vertical lines indicate background fluorescence levels determined from unstained cells. **d**, The groups 1 and 2 were gated as a CCNE2-high CDKN1A-low population and a CCNE2-low CDKN1A-high population, respectively (left, from Fig. 1m), and their forward versus side scatter characteristics (FSC/SSC) were analyzed (right).

**Extended Data Fig. 4.**
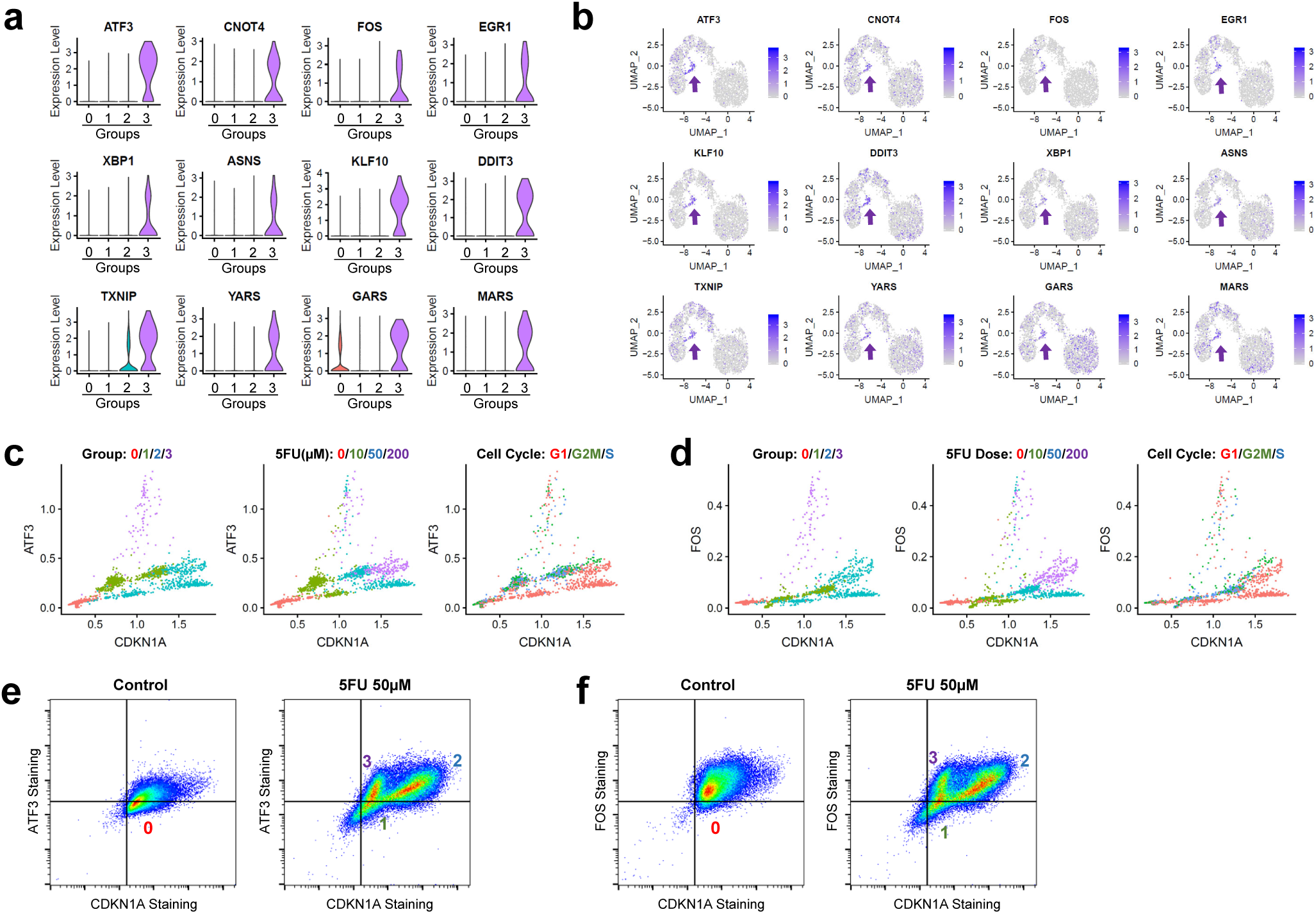
Group 3 was observed in both scRNA-seq and flow cytometry analyses. **a**, Violin plots of the group 3-specific marker genes generated using scaled data without imputation. **b**, Gene expression feature plots of the group 3-specific marker genes. The location of the group 3 is identified by localized expression of these markers, indicated with purple arrows. **c, d**, Scatterplot of imputed gene expression in single cells. Each dot represents data from a single cell colored with its group identity (left) dose of 5FU treatment (center) or estimated cell cycle phase (right). **e, f**, Flow cytometry analysis of indicated protein expression. Cells were subjected to indicated treatments for 24 hours. Horizontal and vertical lines indicate background fluorescence level determined from unstained cells. Areas for the putative groups 0-3 were indicated with corresponding numbers.

**Extended Data Fig. 5.**
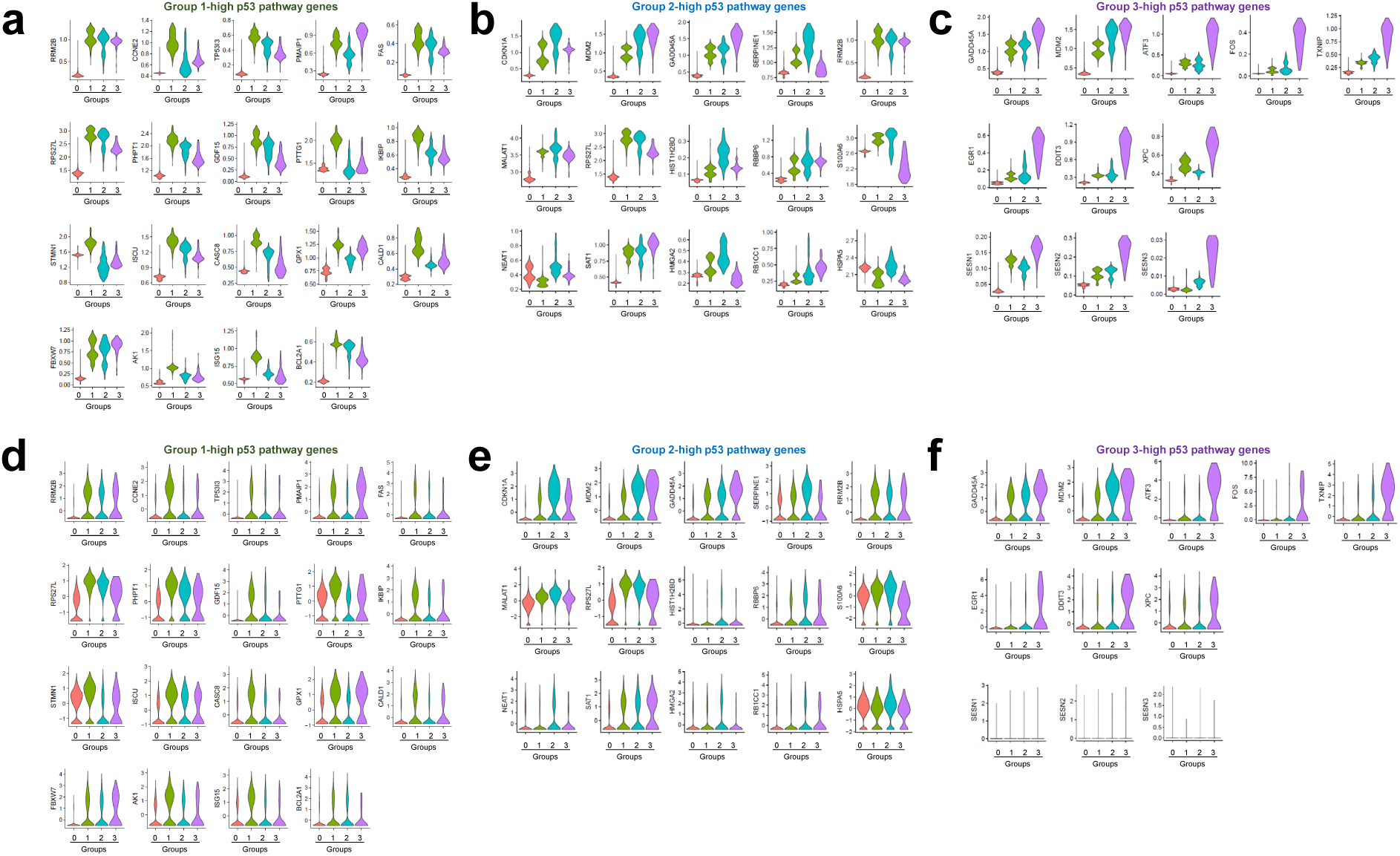
5FU response of p53 pathway genes in different groups of cells. Violin plots of the p53 pathway genes that are in the top 30 markers for the groups 1 (**a, d**), 2 (**b, e**) and 3 (**c, f**), using imputed data (**a-c**) and scaled raw data (**d-f**). *SESN1-3* are not in the top 30 markers but were significantly upregulated in the Stress group.

**Extended Data Fig. 6.**
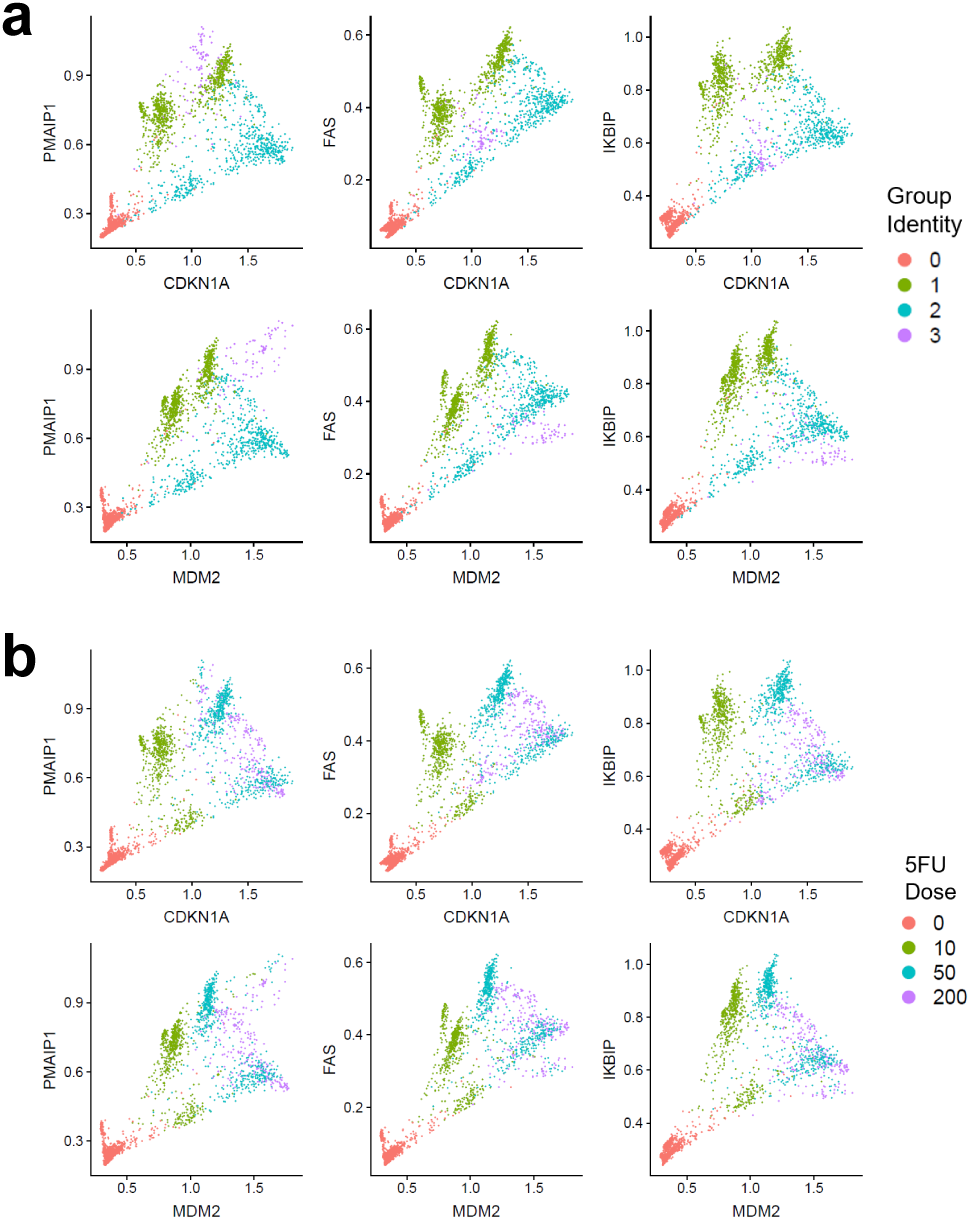
Differential expression of p53 targets according to the group identities of single cells. Scatterplot of indicated gene expression in single cells. Each dot represents individual cell colored with its group identity (**a**) or dose of 5FU treatment (**b**).

## Extended Data Table Legend

**Extended Data Table 1 | Top markers for individual RKO cell groups isolated by clustering analysis.**

Top 30 up-regulated genes for groups 0, 1, 2 and 3: cluster, group identity; gene, gene name; p_val, P value; p_val_adj, adjusted P value; avg_logFC, average fold enrichment. For the Untreated group, only 21 genes appeared significantly upregulated compared to the other groups. For all 5FU-treated groups, all 30 genes were significantly upregulated.

## References

1. Flusberg, D.A., Roux, J., Spencer, S.L. & Sorger, P.K. Cells surviving fractional killing by TRAIL exhibit transient but sustainable resistance and inflammatory phenotypes. Mol Biol Cell 24, 2186–2200 (2013).

2. Holohan, C., Van Schaeybroeck, S., Longley, D.B. & Johnston, P.G. Cancer drug resistance: an evolving paradigm. Nat Rev Cancer 13, 714–726 (2013).

3. Roos, W.P., Thomas, A.D. & Kaina, B. DNA damage and the balance between survival and death in cancer biology. Nat Rev Cancer 16, 20–33 (2016).

4. Jackson, S.P. & Bartek, J. The DNA-damage response in human biology and disease. Nature 461, 1071–1078 (2009).

5. Harper, J.W. & Elledge, S.J. The DNA damage response: ten years after. Mol Cell 28, 739–745 (2007).

6. Derks, K.W., Hoeijmakers, J.H. & Pothof, J. The DNA damage response: the omics era and its impact. DNA Repair (Amst) 19, 214–220 (2014).

7. Macosko, E.Z. et al. Highly Parallel Genome-wide Expression Profiling of Individual Cells Using Nanoliter Droplets. Cell 161, 1202–1214 (2015).

8. Becht, E. et al. Dimensionality reduction for visualizing single-cell data using UMAP. Nat Biotechnol 37, 38–44 (2019).

9. Kho, P.S. et al. p53-regulated transcriptional program associated with genotoxic stress-induced apoptosis. J Biol Chem 279, 21183–21192 (2004).

10. Gene Ontology, C. Gene Ontology Consortium: going forward. Nucleic Acids Res 43, D1049–1056 (2015).

11. Thomas, M.P. et al. Apoptosis Triggers Specific, Rapid, and Global mRNA Decay with 3’ Uridylated Intermediates Degraded by DIS3L2. Cell Rep 11, 1079–1089 (2015).

12. Kanehisa, M. & Goto, S. KEGG: kyoto encyclopedia of genes and genomes. Nucleic Acids Res 28, 27–30 (2000).

13. Park, J.H. et al. Positive feedback regulation of p53 transactivity by DNA damage-induced ISG15 modification. Nat Commun 7, 12513 (2016).

14. Hafner, A., Bulyk, M.L., Jambhekar, A. & Lahav, G. The multiple mechanisms that regulate p53 activity and cell fate. Nat Rev Mol Cell Biol 20, 199–210 (2019).

15. Batchelor, E., Loewer, A. & Lahav, G. The ups and downs of p53: understanding protein dynamics in single cells. Nat Rev Cancer 9, 371–377 (2009).

16. Fischer, M. Census and evaluation of p53 target genes. Oncogene 36, 3943–3956 (2017).

17. van Dijk, D. et al. Recovering Gene Interactions from Single-Cell Data Using Data Diffusion. Cell 174, 716–729 e727 (2018).

18. Tate, J.G. et al. COSMIC: the Catalogue Of Somatic Mutations In Cancer. Nucleic Acids Res 47, D941–D947 (2019).

19. Knight, K. Intracellular Flow Cytometry Staining Protocol.bn dx.doi.org/10.17504/protocols.io.tknekve (2018).

20. Dobin, A. et al. STAR: ultrafast universal RNA-seq aligner. Bioinformatics 29, 15–21 (2013).

21. Genomes Project, C. et al. A global reference for human genetic variation. Nature 526, 68–74 (2015).

22. Stuart, T. et al. Comprehensive integration of single cell data. bioRxiv https://doi.org/10.1101/460147 (2019).

23. Satija, R., Farrell, J.A., Gennert, D., Schier, A.F. & Regev, A. Spatial reconstruction of single-cell gene expression data. Nat Biotechnol 33, 495–502 (2015).

24. van der Maaten, L. & Hinton, G. Visualizing Data using t-SNE. Journal of Machine Learning Research 9, 2579–2605 (2008).

25. Butler, A., Hoffman, P., Smibert, P., Papalexi, E. & Satija, R. Integrating single-cell transcriptomic data across different conditions, technologies, and species. Nat Biotechnol 36, 411–420 (2018).

26. Haghverdi, L., Lun, A.T.L., Morgan, M.D. & Marioni, J.C. Batch effects in single-cell RNA-sequencing data are corrected by matching mutual nearest neighbors. Nat Biotechnol 36, 421–427 (2018).

27. Welch, J. et al. Integrative inference of brain cell similarities and differences from single-cell genomics. bioRxiv, https://doi.org/10.1101/459891 (2019).

28. Nestorowa, S. et al. A single-cell resolution map of mouse hematopoietic stem and progenitor cell differentiation. Blood 128, e20–31 (2016).

29. Yu, G., Wang, L.G., Han, Y. & He, Q.Y. clusterProfiler: an R package for comparing biological themes among gene clusters. OMICS 16, 284–287 (2012).

